# Compensatory traits can explain concave cost function of purely sexual traits

**DOI:** 10.1101/2022.04.27.489663

**Authors:** Masaru Hasegawa

**Affiliations:** Department of Environmental Science, Ishikawa Prefectural University, Nonoichi, Ishikawa 921⍰8836, Japan

**Keywords:** cost-reducing traits, cost of ornamentation, viability selection, sexual selection

## Abstract

The cost of ornamentation is often measured experimentally to study the relative importance of sexual and viability selection for ornamentation, but these experiments can lead to a misleading conclusion when compensatory trait is ignored. For example, a classic experiment on the outermost tail feathers in the barn swallow *Hirundo rustica*, explains that the concave (or U-shaped) aerodynamic performance cost of the outermost tail feathers would be the evolutionary outcome through viability selection for optimal tail length, but this conclusion depends on the assumption that compensatory traits do not cause reduced performance. Using a simple “toy model” experiment, I demonstrated that ornamentation evolved purely though sexual selection can produce a concave cost function under the presence of compensatory traits, which was further reinforced by a simple mathematical model. Therefore, concave cost function (and the low performance of individuals with reduced ornaments) cannot be used to infer the evolutionary force favoring ornamentation, due to a previously overlooked concept, “overcompensation”, which can increase the apparent cost of ornamentation.

## Introduction

Animals sometimes possess ornamentation that seem to have negative effects on survivorship, such as long tails and colorful plumage in birds (Andersson, 1994). Many empirical studies have shown sexual selection for ornaments, explaining the evolution of seemingly costly ornaments (e.g., see Hill & McGraw, 2006 for a review on birds). However, the mere presence of sexual selection does not necessarily mean that sexual selection has been the major selection force causing ornament elaboration, because focal traits can have mainly evolved via viability selection and sexual selection may have caused ornament elaboration only slightly beyond viability optimum (i.e., possible exaptation; Bergstrom & Dugatkin, 2016). Then, it is not surprising to see that researchers have alternatively focused on the cost of ornamentation (e.g., Evans, 1998; see below), because the ornaments’ cost function would clarify the relative importance of sexual and viability selection on the focal traits.

A famous example is those testing aerodynamic performance of long outermost tail feathers in the barn swallows *Hirundo rustica* (Fig. 1; e.g., Evans, 1998; Buchnan & Evans, 2000; Bro-Jørgensen et al., 2007). Although long outermost tail feathers have been shown to be sexually selected (e.g., Møller, 1988; reviewed in Møller, 1994; Turner, 2006; Romano et al., 2016), it remains unclear how and why long tails evolved due to the possible aerodynamic function of long tails (Norberg, 1994). Evans and colleagues have experimentally shortened outermost tail feather length and measured aerodynamic performances of swallows (e.g., Evans, 1998; Buchanan & Evans, 2000; Rowe et al., 2001; also see Evans & Thomas, 1997; Thomas & Rowe, 1997 for the detailed descriptions of predictions). They predicted that, if long outermost tail feathers evolved purely through sexual selection, shortening the length should produce a decrease in aerodynamic cost (Fig. 1 upper left). Or, if it evolved purely through viability selection due to the aerodynamic function, shortening the length should increase the aerodynamic cost (Fig. 1 upper right). Lastly, if viability and sexual selection together favors the evolution of long tails, shortening the tail would decrease the aerodynamic cost first, and then increase the cost beyond the aerodynamic optimum (Fig. 1 upper middle). Their results were consistent with the last prediction with estimated peak values located around 10 mm shorter from the current tail length. Therefore, they concluded that tail feathers mainly evolved through viability selection and sexual selection would have elongated tails only around 10 mm (9–20%) beyond the aerodynamic optimum (and thus many ornithological books follow their conclusions; e.g., Fjeldsa et al., 2020). However, this argument is problematic, because they do not consider compensatory traits (also known as cost-reducing traits), which have been reported to affect aerodynamic cost of long tails in many bird species including barn swallows (e.g., Balmford et al., 1994; Barbosa & Møller, 1995; Møller et al., 1995; Møller, 1996). In fact, several compensatory traits have been reported in barn swallows, including greater wingspans, wing area, aspect ratios, reduced wing loading, and so on (e.g., Møller et al., 1995; reviewed in Husak & Swallow, 2011; also see Moreno & Møller, 1996; Saino et al., 1997; Tubaro, 2003 for other kinds of compensation). Animal performance could be affected by compensatory traits, which is thought to be the case in the peacock *Pavo cristatus*, another model species of sexual selection: peacocks with fully expressed trains, i.e., a costly trait, have a lower metabolic cost of locomotion than those with rudimentary trains possibly due to the presence of compensatory traits (Thavarajah et al., 2016). Similar but less striking results, such as no detectable performance cost of gorgeous ornaments, are repeatedly reported in ornamented animals (e.g., Baumgartner et al., 2011; Trappet et al., 2013; Kojima & Lin, 2019). Assuming costly nature of ornamental traits, these findings are puzzling (although the possibility that all these seeming costly ornaments are virtually cost-free cannot be excluded).

**Figure 1.**
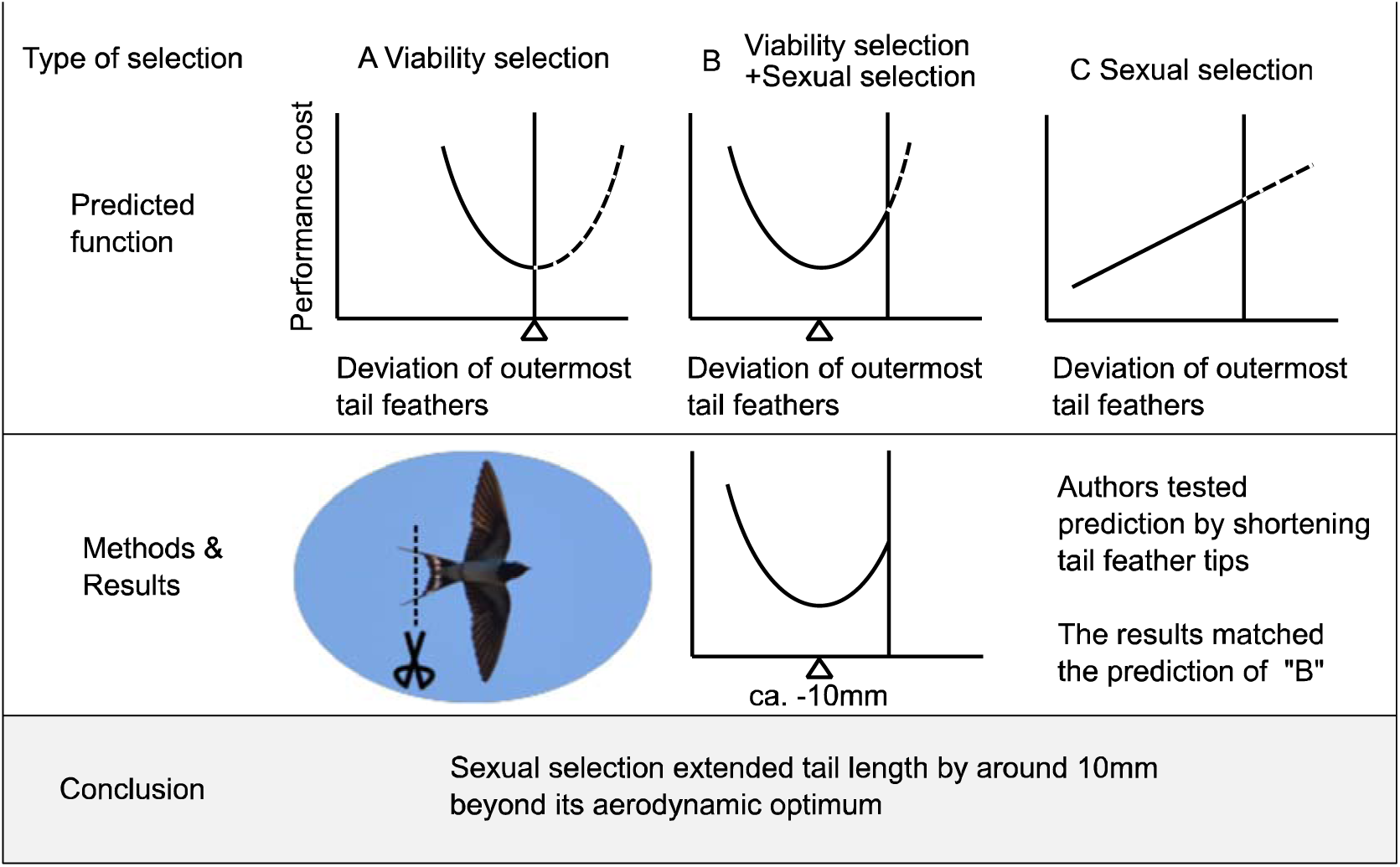
Details of manipulation experiment of Evans and colleagues (see Evans, 1998; Buchanan & Evans, 2000; Rowe et al., 2001 for original figures; also see Evans & Thomas, 1997; Thomas & Rowe, 1997 for similar predictions). The vertical line in each plot indicates current ornament expression, and the triangle indicates ornament expression providing peak performance (note that lower values indicate higher performance here). Although they originally called deviation of outermost tail feathers (or possibly, narrowed part of outermost tail feather length) as a “streamer,” I avoided this wording, because it implies some kinds of aerodynamic function of the ornaments, and because the formal definition of the streamer is unclear (and not to confound size and shape: Matyjasiak et al., 2009). Likewise, I used viability selection rather than natural selection to be precise.

The importance of compensatory traits has repeatedly been advocated (e.g., Møller, 1996; Oufiero & Garland, 2007; Husak et al., 2015). Using a hypothetical data set, Oufiero & Garland (2007) demonstrated that ignoring a compensatory trait led to an incorrect (and opposite) conclusion, at least in a correlational study, because the focal costly trait appeared to have a positive effect on performance if a compensatory trait was not taken into account. Likewise, Husak & Swallow (2011) reviewed compensatory traits and proposed that a simple test of relationships between ornamentation and performance can lead to misleading conclusions (also see Husak et al., 2015 for an updated review). However, although these studies stress that correlational and even manipulation experiments should consider compensatory traits carefully (e.g., Husak & Swallow, 2011), it is still unclear whether and how compensatory traits affect cost function concerning ornament manipulation. Particularly, whether the low performance of individuals with reduced ornaments (and hence concave cost function; see preceding paragraph) can be explained by compensatory traits is not known. As Oufiero & Garland (2007) used a hypothetical data set, a simple model experiment rather than complicated empirical data, would be helpful to demonstrate the potential influence of compensatory traits.

Here, using a simple “toy” model, I examined whether the concave cost function (see Fig. 1) could be produced when the ornament has evolved purely through sexual selection for exaggerated ornamentation with the coevolution of compensatory traits. Although I verbally explained potential confounding effects of compensatory traits above, our simple model experiment would directly demonstrate how compensatory traits affects the performance of (virtual) animals with ornamentation. For this purpose, I used “chuonchuon” (Fig. 2 upper panel), a traditional balancing toy in Vietnam, as a model system. Chuonchuon is a suitable model, because its balance is determined by the whole phenotype, as in the locomotor performance of animals (e.g., see Husak & Swallow, 2011). Different from purely verbal models, simulations, or mathematical models, this kind of simple, physical toy model would be intuitive (and thus suitable) for empirical researchers of animal performance to understand the importance of compensatory traits. Clearly, this toy model is NOT to test the function of particular traits (e.g., swallows’ tails), but to test whether compensatory traits can cause reduced whole-body performance under an experimental reduction of ornamental traits that has evolved purely through sexual selection. Rather than testing the performance of models with arbitrary values of ornamental and compensatory traits, I first determined an evolutionary stable form of chuonchuon based on game theory under the setting that long tails have been sexually selected (i.e., providing reproductive advantages) and its negative effects on balance (i.e., survival of chuonchuon) could be reduced by a compensatory trait, extended wings. Then, I tested the cost function of ornamentation by experimentally shortening tail length, as in the previous manipulation experiments in swallows listed above. To test the generality of the finding of chuonchuon, I also provided a simple (or even rudimentary) mathematical model. Even such a simple model would be useful to determine whether and, if any, in which condition the apparent concave cost function of ornament can be found. I discussed the evolutionary implications of the observed pattern.

**Figure 2.**
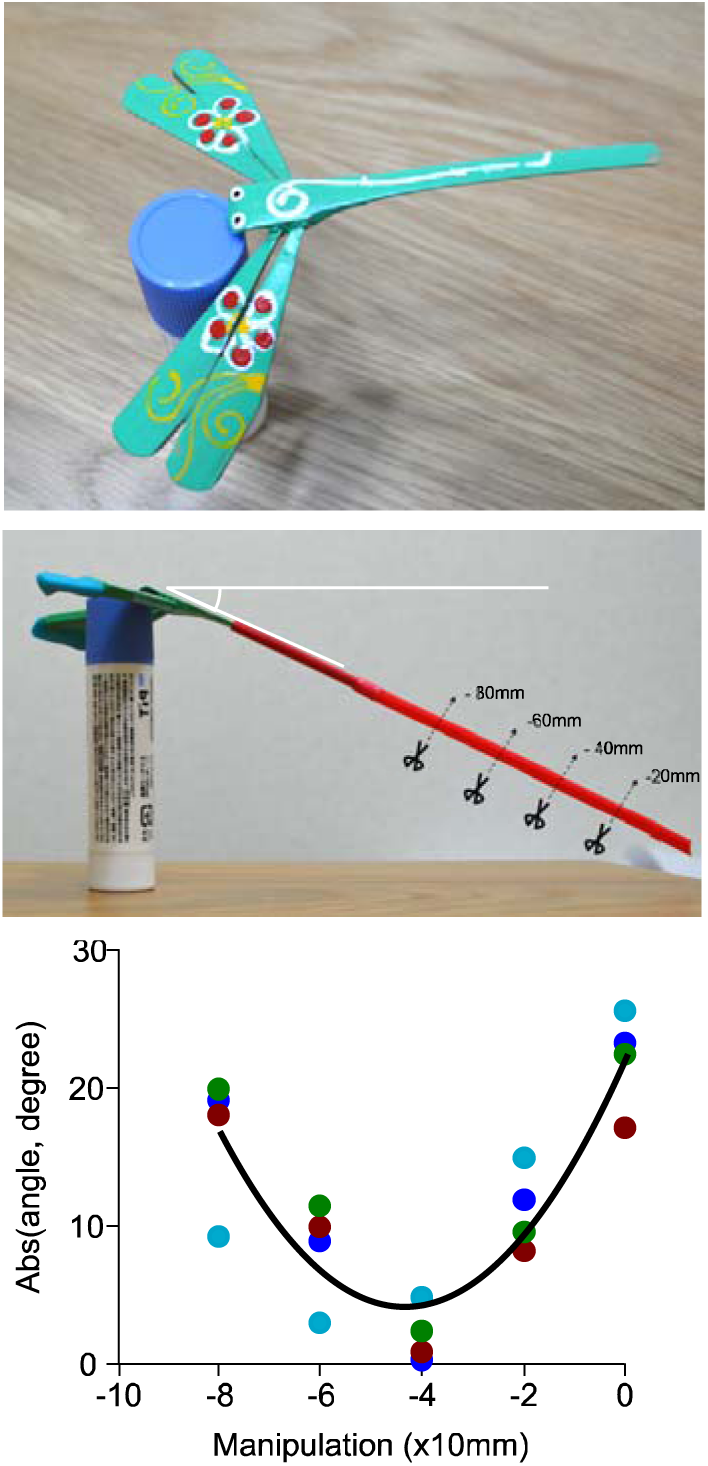
Chuonchuon (upper panel), chuonchuon with elongated tails and extended wings (middle panel), showing how manipulation was conducted, and results of chuonchuon experiment (lower panel). A simple quadratic regression line is denoted (see Table 1 for a formal analysis). Individual identity was denoted by different coloration. Mean ± SE values of stability, measured by absolute angle deviated from horizontal plane, in ancestral state shown in upper panel was 4.87 ± 1.78 (range: 1.38–8.87). Not surprisingly, tail elongation without extended wings (i.e., models without compensatory traits) made models unbalanced in the range of the current study (e.g., when elongated tails = 30 mm, which corresponds to the manipulation = −80 mm, mean stability ± SE = 36.60 ± 2.30, range = 32.69–42.93).

**Table 1.**
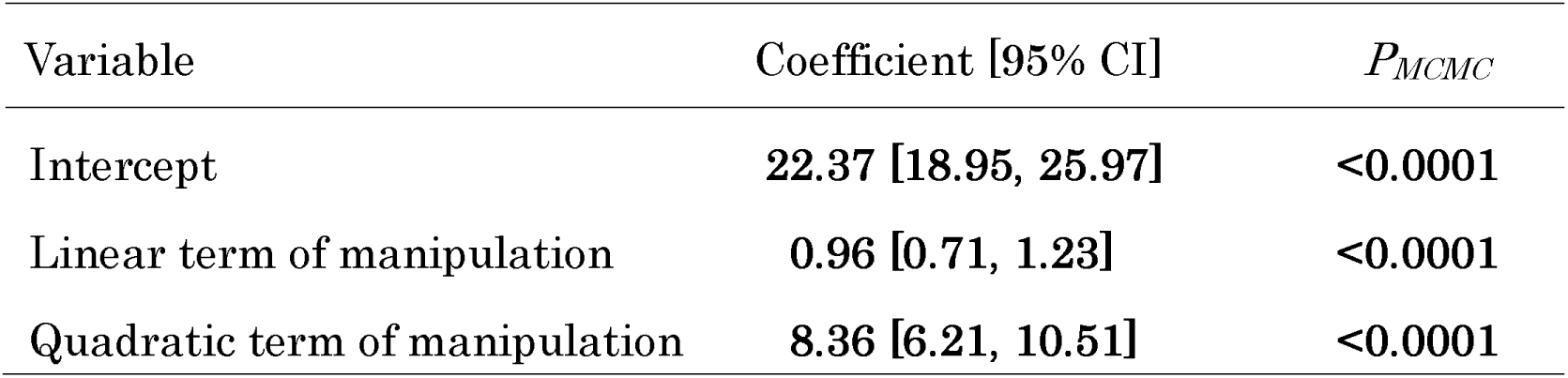
Bayesian linear mixed-effects model of stability of chuonchuon in relation to experimental manipulation of elongated tails (*N_ID_* = 4, *Ntotal* = 20). The dependent variable is the absolute angle of chuonchuon deviated from the horizontal plane. I included chuonchuon ID as a random factor (variance = 0.20) in addition to units (variance = 15.39). Significant test results (i.e. *P* < 0.05) are indicated in bold

## Material and methods

### Experimental setup

I used four commercially available “chuonchuon” for my experiment (ca. 7 cm length: Tirakita, Japan). I used unmanipulated chuonchuon as an ancestral state, in which neither sexually selected ornamentation nor compensatory traits have yet been evolved. Then, I used a plastic straw (0.4 g, 4 mm width and 160 mm length; Strix design, Japan) as an elongated tail. As a compensatory trait, I made extended wings using clay (0.8 g per wing, i.e., twice as heavy as a plastic straw to account for the lever principle; Fig. 2 middle panel).

I here simply set the situation with a set of dichotomous events, that is, only balanced chuonchuon can “survive,” and only longer tailed chuonchuon in the population can “reproduce.” Then, the ancestral form can survive anytime, but the tail-elongated form cannot survive when elongated tails make them unbalanced (I used a cut-off point as 30° deviation from the horizontal plane, here). Tail elongation made models unbalanced in the range of the current study (i.e., maximum length to −80 mm shortened tails; see Fig. 1) and thus they cannot survive without compensatory traits, i.e., extended wings. Even when it survived, the ancestral form cannot have descendants when any derived forms with elongated tails (i.e., those with longer tails than ancestral form) exist. Because my interest here is not in the actual evolutionary trajectory but the cost function of the derived form, I obtained the evolutionary stable form by comparing the “fitness” of each form. For simplicity, I assumed continuous variation in tail length but assumed two discrete states of the extended wings (i.e., with and without extended wings). In this case, the evolutionary stable form is a chuonchuon with maximum elongated tails with extended wings. This is because chuonchuon with maximum elongated tails and extended wings can survive (i.e., balanced) and thus reproduce under any circumstances (i.e., they have the longest tails compared to all other forms). I did not subdivide compensatory traits (e.g., small/medium/large), because the purpose of the current study is not to quantify the exact peak performance or identify the best size of compensatory traits but to examine the effect of compensatory traits.

### Measuring performance

As in manipulation experiments in the barn swallow (Buchanan & Evans, 2000; Rowe et al., 2001), I shortened the extended tails of the evolutionary stable form (i.e., a chuonchuon with maximum elongated tails as ornamentation with extended wings as a compensatory trait; see above) of chuonchuon models by 0 mm, 20 mm, 40 mm, 60 mm, and 80 mm, and took two pictures each time. From each of the two pictures per individual chuonchuon, I measured the absolute angle of chuonchuon deviated from the horizontal plane as stability (or balance-ability: Fig. 2 middle panel) using ImageJ software. Then, I averaged the two measurements (repeatability = 0.99, F_19,20_ = 270.24, *P* < 0.0001; Lessells & Boag, 1987) to have a representative estimate of stability for each treatment (i.e., tail shortened by 0 mm/20 mm/40 mm/60 mm/80 mm) in each chuoncuhon.

### Statistics

To account for individual variation in performance, I used a Bayesian linear mixed-effects model with a normal error distribution to examine the stability of chuonchuon given tail shortening. Here, stability, measured as the absolute angle of chuonchuon (see above), was used as a response variable. Individual identity was used as a random effect. By calculating the Brooks-Gelman-Rubin statistic (Rhat), which must be <1.2 for all parameters (Kass et al., 1998), the reproducibility of the MCMC simulation was confirmed. Data analyses were conducted using the R statistical package (ver. 4.1.0; R Core Team, 2021), using the function “MCMCglmm” (with its default setting) in the package “MCMCglmm” (Hadfield, 2010).

### Mathematical model

To test whether a quadratic cost function of purely sexual traits (see above) can be found in the presence of compensatory traits, I used a simple mathematical model. I used here an equation for a bivariate quadratic selection surface, characterized by slope and curvilinear terms, as commonly used in evolutionary biology (cf. Lande & Arnold, 1983; Arnold et al., 2001; Arnold, 2003; Stinchcombe et al., 2008; see Results). This selection surface can be used to approximate complex empirical performance surfaces regardless of the shape of the actual performance surface (Arnold, 2003). I used the same symbols for each term as those used in the literature on selection surface (e.g., 0.5*γ_ii_, instead of q_ii_, though the latter is often used in multiple regression models: Stinchcombe et al., 2008).

## Results

### Toy model

I found a significant quadratic relationship between tail length manipulation and stability of the chuonchuon (Table 1; Fig. 2 lower panel): the stability of the chuonchuon, measured as an absolute angle deviated from the horizontal plane, increased with decreasing tail length until the peak value where chuonchuon had the maximum estimated stability and further tail shortening beyond the peak value decreased stability.

### Mathematical model

Suppose that animal performance can be approximated using the following equation (cf. Lande & Arnold, 1983; Arnold et al., 2001; Arnold, 2003; Stinchcombe et al., 2008):

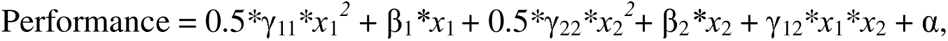

where *x*_1_ denotes the expression of the ornament, *x*_2_ denotes the expression of compensatory trait(s), and γ_11_, γ_22_, γ_12_, β_1_, and β_2_ are coefficients for each variable with α as a constant.

For simplicity, imagine an elliptical performance surface where the optimal performance can be set at (*x*_1_*,x*_2_) = (0,0), so that any additions of the ornament and compensatory traits decrease performance, which is accomplished by the condition, β_1_ = β_2_ = 0, with γ_11_ < 0, γ_22_ < 0, and γ_12_^2^ < γ_11_*γ_22_ (note that, when γ_12_^2^ > γ_11_*γ_22_, there will be a saddle in the performance surface: Stinchcombe et al. 2008). Assuming that the performance surface has a hill with an upward tilting axis (i.e., for each ornament expression, the decline of performance can be minimized by the presence of certain amounts of compensatory traits, which is accomplished by γ_12_ > 0; see Fig. 3 for an example), the above equation can be reduced as follows:

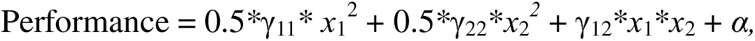

**Figure 3.**
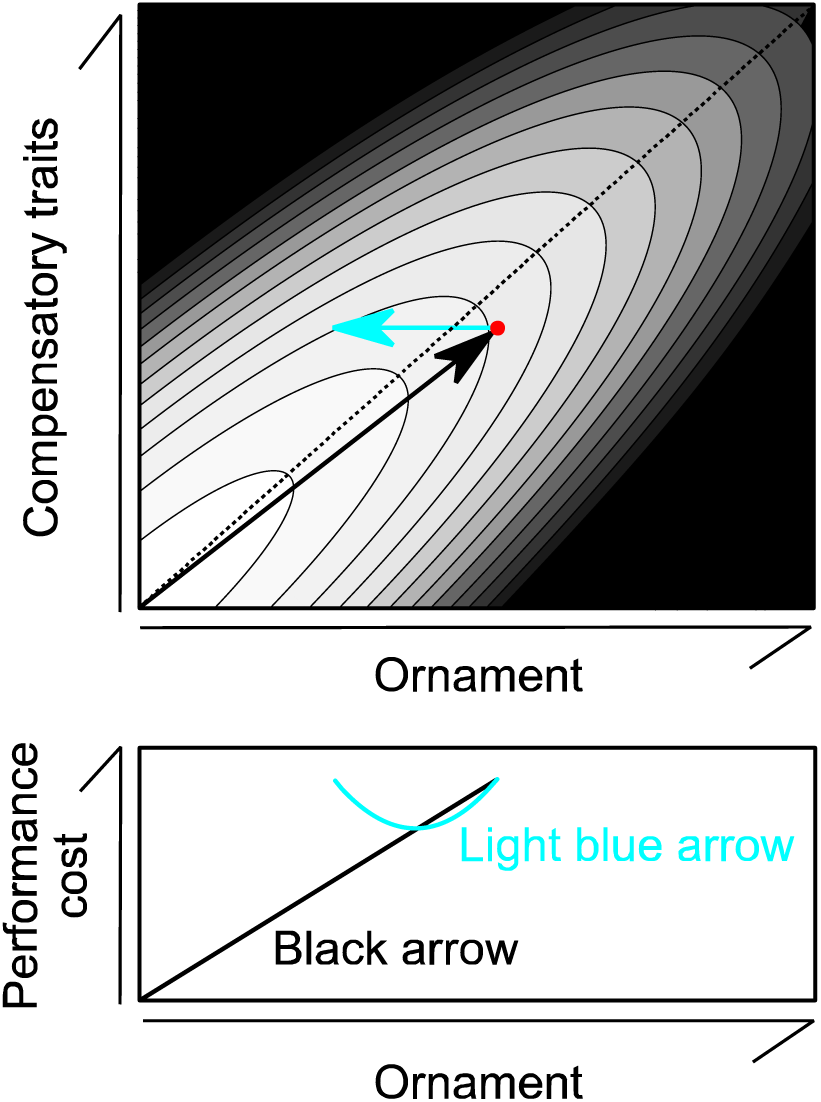
An example of hypothetical performance surface in relation to ornament and phenotypic expression of compensatory traits (upper panel). Darker background coloration indicates lower performance (note that the best performance is attained in the lower left corner). Black arrow indicates hypothetical evolutionary pathway to the current state (filled circle). Light blue arrow indicates phenotypic change when researchers experimentally reduced ornamentation (in which compensatory traits remain unchanged). Lower panel shows performance cost measurement (which is here denoted as the best performance minus the performance of the focal coordinate) in relation to ornament expression along with black or light blue arrows. The evolutionary pathway to the current state is biased toward right side because of intense sexual selection favoring long tails (i.e., actual evolutionary pathway depends on sexual selection in addition to viability selection due to performance cost and other costs, such as production costs of ornament and compensatory traits; see text). Narrow contour width indicates that small deviation from integrated, co-opted trait sets had a strong negative effect on performance, as predicted by Norberg (1994). Here, for the illustrative purpose, I assumed that performance is determined by –4*x*^2^ – 4*y*^2^ + 7*xy* where *x* denotes ornament expression whereas *y* denotes the expression of compensatory traits. The current state is determined here as the point in which the sum of performance surface and sexual selection surface (here, 10*x*) is maximized (see text for detailed explanation). `Dotted line is shown here as a ridge line`.

with the condition, γ_11_ < 0, γ_22_ < 0, γ_12_^2^ < γ_11_*γ_22_, and γ_12_ > 0. In this bivariate performance surface, the experimental manipulation of the ornament is equivalent to changing the *x*_1_ value while *x*_2_ value remains unchanged from the current value of compensatory traits (say, k; note that k > 0 under the presence of compensatory traits). Then, by substituting *y* = k into the above equation, the quadratic function of *x*_1_ is obtained:

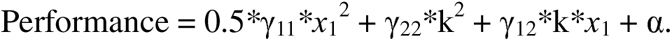

This can be re-written as follows:

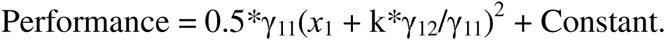

Now, it is clear that we can have a convex performance function (and hence, concave cost function) of ornament, because k*γ_12_/γ_11_ < 0 (note that k > 0, γ_12_ > 0, and γ_11_ < 0; see above). Given that sexual selection favors the exaggeration of the ornament (i.e., the bivariate sexual selection surface can be an increasing function of *x*_1_), the current value of the ornament should be located on the right side of the ridge line (i.e., beyond the upward tilting axis of the hill; see above; also see Fig. 3) where sexually selected advantages offset low performance of ornamented animals. When experimentally reducing the ornament expression, we find that performance increases until the ornament is reduced to be k*γ_12_/γ_11_, and then decreases (Fig. 3).

Retrospectively, this equation indicates that an incremental cost function of sexual traits, which is supposed to be common under manipulative experiments of the ornament (see Fig. 1C), can be predicted only when k*γ_12_/γ_11_ ≥ 0. This can be accomplished when k = 0 (i.e., when compensatory traits are absent for some reason, such as lack of genetic variation) or when γ_12_ ≤ 0 (i.e., when seemingly compensatory traits are not actually compensatory traits but just costly traits like ornament with their joint costs being additive or multiplicative). Although this condition is confined to an elliptical performance surface as assumed above (i.e., when γ_12_^2^ > γ_11_*γ_22_), this is a reasonable assumption. When there is a saddle in the performance surface (e.g., see Fig. S1 left panel), there is no optimal performance different from elliptical performance surface. Furthermore, it accompanies the changing function of the ornament, in which ornament is not a purely sexual trait but can be a functional trait in some ranges of variables, which is beyond the scope of the current study (because here we focused on purely sexual traits; see Introduction). Under the condition γ_12_^2^ = γ_11_*γ_22_, the cost of the ornament is perfectly compensated by compensatory traits, and thus animals can have any ornament value together with the corresponding compensatory traits without performance costs (Fig. S1 right panel). In other words, the cost of ornament can be virtually absent under this condition. As in elliptical performance surfaces (see above), performance increases until the ornament is reduced to k*γ_12_/γ_11_, and then decreases in this case.

## Discussion

The main finding from the chuonchuon experiment is that the apparent concave cost function can evolve purely through sexual selection in the presence of compensatory traits (i.e., with no viability advantage of ornamentation), which is further reinforced by a mathematical model. The importance of compensatory traits has repeatedly been advocated (reviewed in Møller, 1996; Husak & Swallow, 2010; Husak et al., 2015), but previous studies assume that compensatory traits still function as “cost-reducing” traits regardless of the situation (e.g., see Husak et al., 2015, p. 18). Here, I showed that “overcompensation” could decrease performance, producing a concave cost function, which is not surprising as individual performance is determined not by single traits but by the whole phenotype. In the current toy model, extended wings as a compensatory trait worsened the whole-body performance measured as stability, once the part of the elongated tail was removed. As an old Chinese proverb say, “too much is as bad as too little” at least in some cases. The assumption that the cost function can reveal the relative importance of sexual and viability selection (e.g., Buchanan & Evans, 2000) is therefore unreliable. In other words, the prediction of sexual selection (i.e., Fig. 1C) is no longer valid and can be replaced by a concave cost function (as in Fig. 1B) at least under the presence of compensatory traits, indicating that the presence/absence of viability selection is undetectable (see Fig. S2 for the corrected prediction).

Of course, the current experiment using chuonchuon models does not mean that sexual selection always produces a concave cost function (and does not mean that the same physical property, i.e., stability, determined the evolution of ornamentation, such as swallows’ tail, and compensatory traits). The actual compensatory traits would include multiple traits (see Introduction), and hence the current balancing toy model should be regarded as a “basic” model possessing single compensatory trait (or, more accurately, a set of compensatory traits, since extended wings in fact alter multiple dimensions of the wing morphology). Still, even without mathematical model (see results), it seems likely that overcompensation will often reduce locomotor performance, particularly when locomotion requires co-opted, integrated phenotype (e.g., aerial locomotion). In fact, concerning the function of long tails in barn swallows, Norberg (1994) mentioned that “*any experimental shortening and lengthening of the outer tail feathers is likely to upset an original co-adapted character set*…” (p. 231). When compensatory traits co-evolved with the sexually selected tail length to minimize the aerodynamic cost of tail length (Møller, 1996), an experimental reduction in tail length alone cannot clarify the aerodynamic cost of tail length, because the manipulation did not follow the actual evolutionary pathway (see Fig. 3; also see Fig. S3 for an alternative scenario in which ornament has some aerodynamic function). Also, all macroevolutionary studies of these aerial insectivores so far have supported sexual (rather than viability) selection explanation (e.g., Hasegawa et al., 2016; Hasegawa & Arai, 2017, 2018, 2020a,b, 2021, 2022), demanding further experimental studies, possibly using a smaller amount of manipulation (so that overcompensation can be avoided) in many species with various ornamentation (i.e., phylogenetic comparative “experiment”; Fig. S4). An alternative approach, the manipulation of compensatory traits (sensu Møller, 1996; see also Fig. 3) would in theory be valuable to show the effect of overcompensation, but is impractical due to the multi-dimensionality of compensatory traits (e.g., manipulations of wing size and shape are often difficult to conduct).

In addition, the current experiment provides useful insights. First, the cost of ornamentation can be transformed from one type to another. Extended wings enhanced the stability of chuonchuon with elongated tails, though they required additional investment (i.e., a production cost) instead (also see Møller, 1996 for a similar argument). Furthermore, the production cost of compensatory traits can be much higher than the production cost of the ornament itself (see Methods). It is straightforward to conclude that a single measure of cost (e.g., aerodynamic cost) is inappropriate to approximate the total cost (and thus viability disadvantage) of ornamentation, even when the production cost of ornamentation itself seems negligible at first glance, as is the case for the outermost tail feathers in barn swallows (Fig. 1). Although we discussed here about a simple compensatory trait (i.e., extended wings) with a production cost, alternative, but not mutually exclusive, kinds of costs can be involved: for example, by changing the position of pivot point via the change of the body shape in the toy model (i.e., possible developmental cost). Such morphological changes will accompany several physiological and behavioral changes (and thus may also require some form of maintenance cost) in living animals. We therefore should keep in mind that any, or at least many, kinds of costs can be involved to compensate costly ornaments, and that the considerations of all these costs might be impractical.

In summary, the current model experiment demonstrated that a concave cost function, which is often thought to result from viability advantage of focal ornaments, can be observed when sexual selection has favored the evolution of the costly ornaments, due to the presence of compensatory traits. Given that compensatory traits coevolved with sexual traits (Møller, 1996), the manipulation of sexual traits alone would not clarify the evolutionary force on sexual traits (see Figs. 3 & S3). Future studies should consider that overcompensation can be detrimental and that more sophisticated experimental design is needed when inferring selection pressure on (and the evolutionary history of) ornamentation.

## Acknowledgments

I thank Dr. Emi Arai, Dr. Shumpei Kitamura and his lab members at Ishikawa Prefectural University for their kindest advices and supports. I also thank Prof. Michael D. Jennions, reviewers, and English-proofing company (Enago) for their helpful comments. MH was supported by the KAKENHI grant of the Japan Society for the Promotion of Science (JSPS, 19K06850). I am grateful to valuable comments by Dr Alex McQueen and anonymous reviewers, which greatly improve the quality of the manuscript.

## Data Availability Statement

Data attached as Table S1 will be deposited into osf.io once accepted.

## Compliance with ethical standards

Conflict if interests The author declares no conflict of interest.

Ethical approval This study does not include any treatments of animals, as all the information was gathered from model experiments.

## Author Contribution

I performed data analysis and wrote the manuscript alone.

---

**Figure S1.**
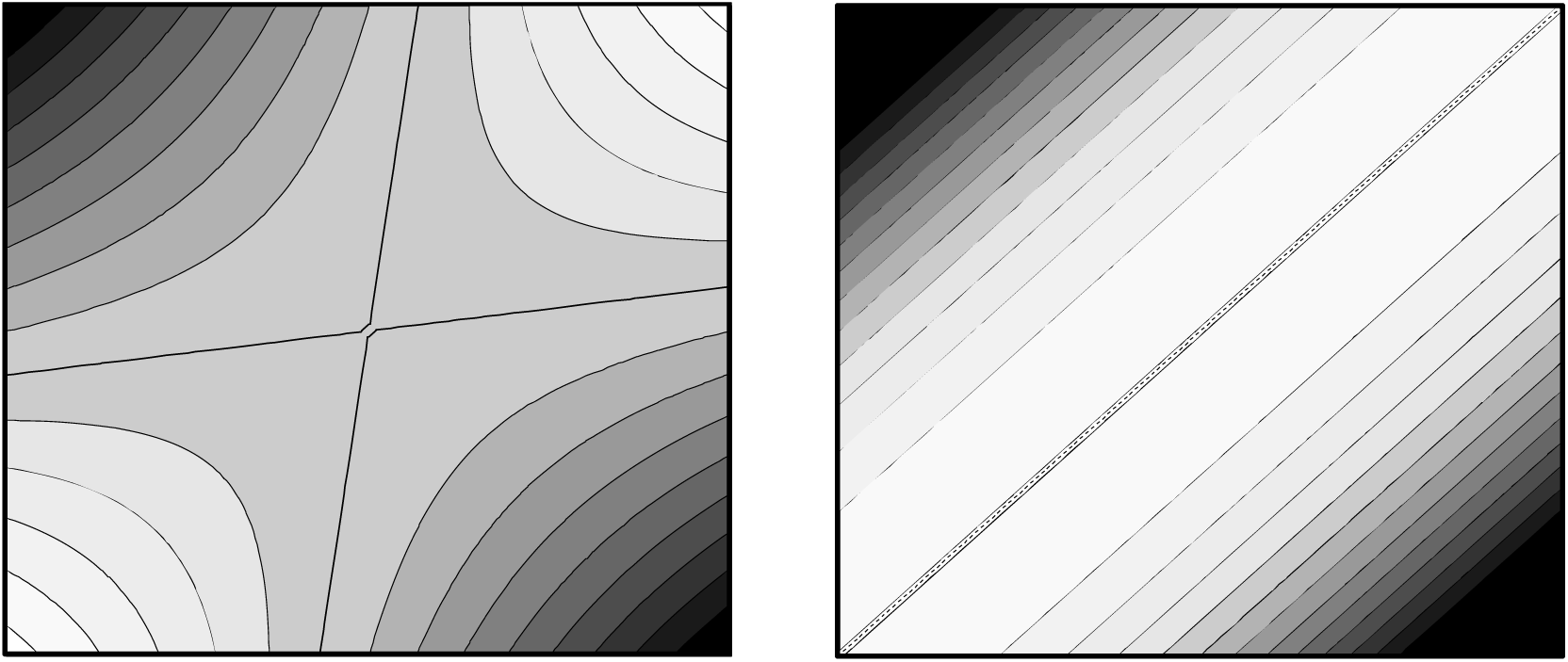
An example of performance surface with a saddle (left panel) and an example of performance surface with a ridge (right panel), both of which are given by performance surface = 0.5*γ_11_\**x_1_^2^* + β_1_*\*x_1_* + 0.5*γ_22_\**x_2_^2^* + β_2_*\*x_2_*+ γ_12_\**x_1_*\**x_2_* + α with the condition, γ_12_^2^ > γ_11_*γ_22_(former) and γ_12_^2^ = γ_11_*γ_22_ (latter). Darker background coloration indicates lower performance.

**Figure S2.**
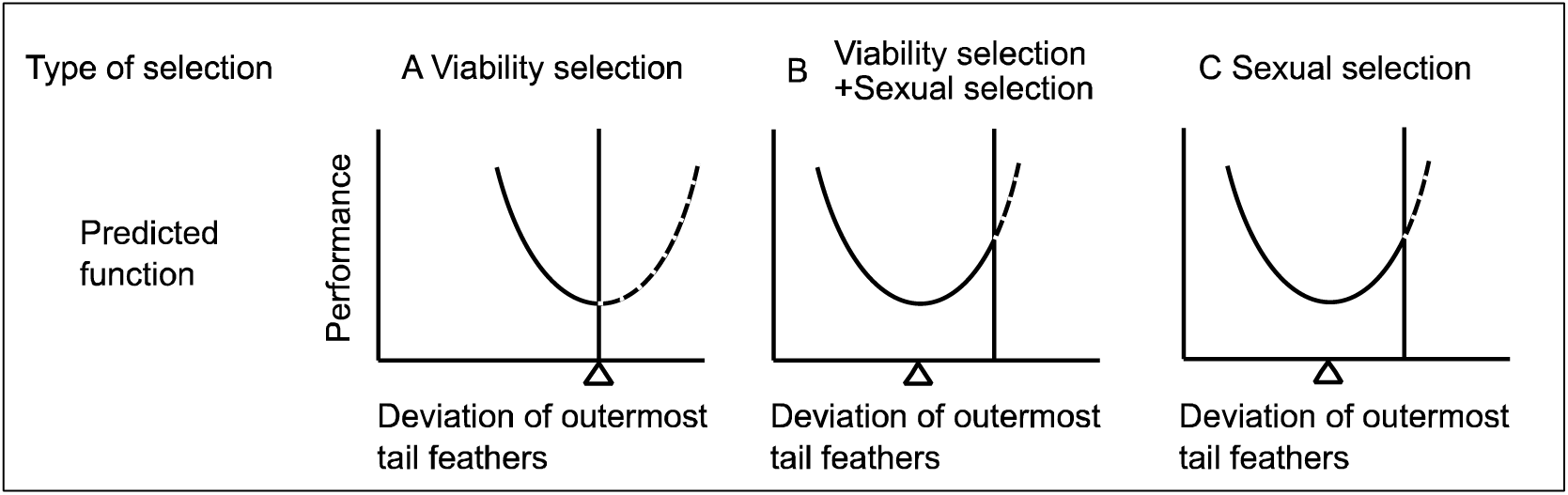
The “corrected” predictions for manipulation experiment of Evans and colleagues under the presence of compensatory traits. Note that two type of scenarios (B and C) cannot be distinguished. See text and Fig. 1 for detailed explanations (also see Figs. 3 & S3).

**Figure S3.**
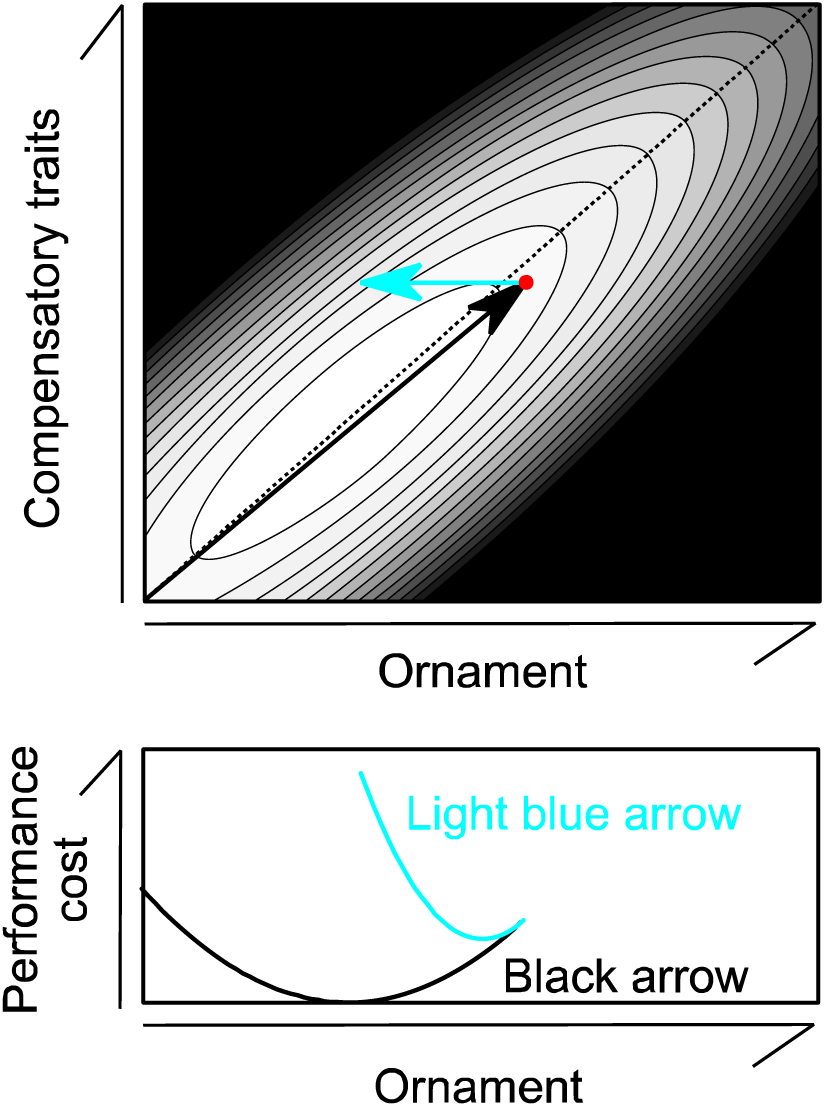
An example of hypothetical performance surface in relation to ornament and phenotypic expression of compensatory traits (upper panel). Darker background coloration indicates lower performance (note that the lightest background located near the current state, shown as a filled circle, in this example). Black arrow indicates hypothetical evolutionary pathway to the current state. Light blue arrow indicates phenotypic change when researchers experimentally reduced ornament expression (in which compensatory traits remain unchanged). Lower panel shows performance cost measurement (which is here denoted as the best performance minus the performance of the focal coordinate) in relation to ornament expression along with black or light blue arrows. The evolutionary pathway to the current state is biased toward right side because of intense sexual selection favoring long tails (i.e., actual evolutionary pathway depends on sexual selection in addition to viability selection due to flight cost and other costs, such as production costs of tail length and compensatory traits; see text). Narrow contour width indicates that small deviation from integrated, co-opted character sets had a strong negative effect on flight performance, as predicted by Norberg (1994). Here, for the illustrative purpose, I assumed that performance is determined by –4(*x* – 3)^2^ – 4(*y* – 3)^2^ + 7(*x* – 3)*(*y* – 3) where *x* denotes ornament expression whereas *y* denotes the expression of compensatory traits. The current state is determined here as the point in which the sum of performance surface and sexual selection surface (here, 5*x*) is maximized (see text for detailed explanation).

**Figure S4.**
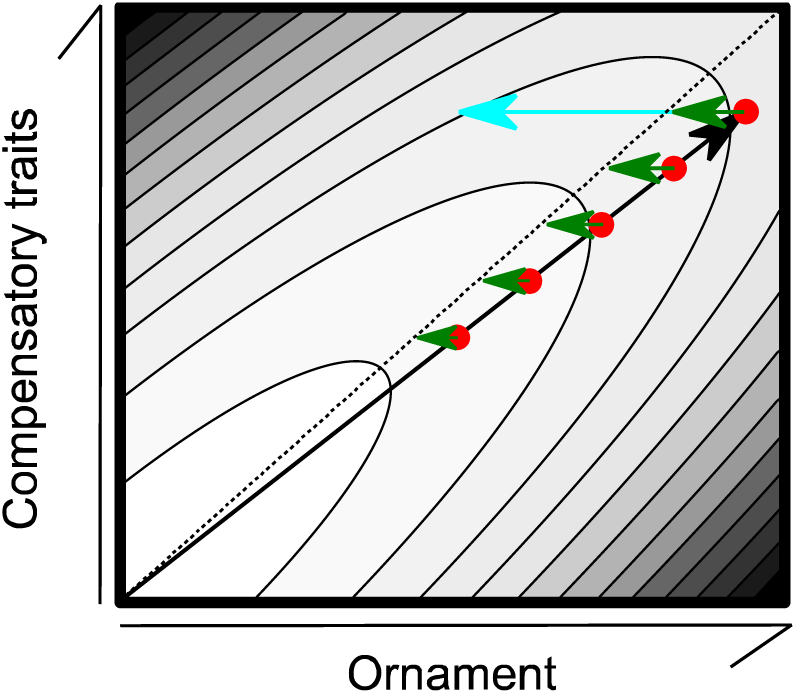
A phylogenetic comparative experiment. Rather than reducing a large extent of ornamentation in a single species, which can result in overcompensation (light blue arrow), researchers should reduce small extents of ornamentation in multiple species with varying expressions of ornamentation, without being bothered by overcompensation (dark green arrow). Black arrow indicates a hypothetical evolutionary pathway to the current state of the most ornamented species. Less-ornamented species are assumed to be present somewhere near the evolutionary pathway (i.e., they are here assumed to be hypothetical ancestral states of the most ornamented species; sensu Matyjasiak et al. 2009). See Fig. 3 for detailed information (note that this figure is derived from the upper panel in Fig. 3). For easy understanding, dotted line is shown here as a ridge line, above which overcompensation occurs.

